# Biofilm of *Klebsiella pneumoniae* minimize phagocytosis and cytokine expression by macrophage cell line

**DOI:** 10.1101/623546

**Authors:** Sudarshan Singh Rathore, Lalitha Cheepurupalli, Jaya Gangwar, Thiagarajan Raman, Jayapradha Ramakrishnan

## Abstract

Infectious bacteria in biofilm mode are involved in many of persistent infections. Owing to its importance in clinical settings many *in vitro* and *in vivo* studies have analysed the structural and functional properties of biofilm, its resistance to antibiotic exposure etc. Currently the immune mechanism toward the clearance of biofilm infections is being investigated. *K. pneumoniae* is one of the major leading causes of biofilm infections on indwelling medical devices. There was no previous literature that demonstrates the interactions of macrophage cells lines and *Klebsiella* biofilm, as the first report, we investigated the *in vitro* response of *Klebsiella* biofilm to phagocytosis and cytokine expression. We developed an *in vitro* model to study the interactions of *Kebsiella* biofilm and macrophage. The phagocytosis assay was performed for heat inactivated and live biofilm. A similar phagocytic response against both biofilms were observed when these cells were exposed to RAW 264.7 macrophages. Also, the expressions of TLR2, iNOS, inflammatory cytokines such as IL-β1, IFN-γ, IL-6, IL-12, IL-4, TNF-α and anti-inflammatory cytokines, IL-10 during phagocytosis were analysed. These results collectively demonstrated that the rate of phagocytosis was an average of 15% for both biofilms. Also, when activated macrophage was exposed to heat-inactivated or live biofilms, there was a significant increase in proinflammatory cytokine genes together with expected increase in TLR2 and iNOS. Thus, it is clear that macrophage response against biofilm producing *K. pneumoniae* results in increase in phagocytic rate and a corresponding increase in inflammatory cytokine gene expression which could be important for clearing *K. pneumoniae* cells.

## Introduction

*K. pneumoniae* is a Gram-negative, encapsulated opportunistic pathogen that colonizes almost every part of the human body with most preferred site being the respiratory, gastrointestinal and urinary tracts [1]. *K. pneumoniae* causes both hospital and community-acquired infections [2]. Pneumonia, meningitis, urinary tract infections and catheter-related bloodstream infections are the potential illness caused by this bacterium [3]. The major risk factors associated with *K. pneumoniae* infection includes central venous catheterization, urinary catheterization, mechanical ventilation, prolonged stay in intensive-care unit, low birth weight in preterm infants and individuals with impaired immunity [4].

*Klebsiella* spp are characterized by the presence of capsular polysaccharides (CPS), type 1 and 3 fimbriae as the major virulence factors. These cellular components play an important role in the adhesion and colonization of host tissues. In addition, these virulence factors are essential for biofilm formation on indwelling medical devices and persistent infections [2].

In order to overcome the infections caused by planktonic and biofilm of *K. pneumoniae*, both humoral and cell-mediated immune defenses are involved [5]. The host immune responses to planktonic *K. pneumoniae* have been extensively investigated [6,7], however, information related to host immune responses to *K. pneumoniae* biofilms remains to be explored The role of immune defenses and the immune evasion mechanism by *Klebsiella* sp have been studied in both *in vitro* and *in vivo* models [8]. The first line defense mechanism includes ciliary lining, the flow of urine, peristalsis, mucus, bile and digestive enzymes [9]. Once it overcomes these mechanical barriers, humoral and cellular innate defenses functions to eliminate the pathogen. Humoral defenses consist of, i. complement system, which forms membrane attack complex and cause cell lysis, also it act as opsonin and enhances phagocytosis ii. defensins, which are bactericidal and disrupt cell membrane iii. transferin that sequester iron and facilitate bacterial growth. However, most *Klebsiella* strains appear to evade or resist the complement-mediated membrane attack lysis and opsonophagocytosis in both *in vitro* and *in vivo* models [5].

In addition there has also been studies to show the importance of IL-8, CXCL1, and leukotriene B4 during infection [5,10]. Apart from macrophage, neutrophils are found to have greater phagocytic activity than alveolar macrophages in clearance of *Klebsiella* infection. Similarly, the functions of TLR during *Klebsiella* infection have been highlighted. Mice with defective TLR4 signaling were observed to show increased bacterial load and consequent mortality, suggesting TLR4 importance [6]. Upon bacterial recognition by pattern recognition receptors such as TLR4, it initiates the pulmonary innate immunity against *Klebsiella* by the release of cytokines such as TNF-α. Another important cytokine in defense against *K. pneumoniae* is IL-12, which is essential for the induction of IFN-γ. IFN-γ is a critical mediator that involves in bacterial clearance and prevention of dissemination from the lungs and improved survival rates in case of localized pulmonary *Klebsiella* infection [7] and the host responses towards systemic infection with the same bacteria appeared to be IFN-γ independent. On the contrary the detrimental role of anti-inflammatory IL-10 in persistent *K. pneumoniae* infection has also been well proved in studies [11]. However most of these studies have looked at planktonic cells and *K. pneumoniae* biofilm interaction with immune cells is not really clear.

*K. pneumoniae* is capable of causing biofilm infections in tissues of the susceptible host as well as on indwelling medical devices such as catheters and prostheses. The establishment of biofilm on the tissues of the susceptible host facilitate the expression of virulent traits and causes infections of upper and lower respiratory tract, otitis media, tonsillitis, cystic fibrosis, urinary tract infections, chronic bacterial prostatitis [12]. Several studies in the last decade had proven the increased antibiotic resistance of *Klebsiella* in biofilm state than planktonic cells [13,14]. Biofilms have also been hypothesized to contribute to reduced bacterial clearance by the innate defense mechanisms of the host.

This includes limited penetration of leucocytes and their antimicrobial products into the biofilm, reduced phagocytic activity, and modified gene expression patterns and genetic shifts. In a first attempt we tried to understand the interactions between macrophage cells and *K. pneumonaie* biofilms using *in vitro* model in terms of phagocytic activity, TLR, iNOS, proinflammatory and anti-inflammatory cytokine expression during phagocytosis.

## Materials and Methods

### Strain

The study strain *Klebsiella pneumoniae* NDM-05-506 was procured from Microbial Culture Collection Centre at Pune, India, which was already proven by our lab to be a proficient biofilm producer [15].

### Cell lines

Raw264.7 macrophage cell line was procured from National Centre for Cell Science, Pune, India. Cells were maintained in DMEM with 10% FBS and 1X antibiotics in 5% CO_2_ with 95% moisture at 37°C.

### Preparation of biofilm for interaction study

To study the interactions of macrophage and *Klebsiella* biofilm, the bacteria was cultured overnight in nutrient broth at 37°C. The culture was then centrifuged at 5,000 X g for 10 minutes. The cell pellets was washed with PBS and re-suspended in nutrient broth. The bacterial cells were plated at a density of 10^6^ cells/mL on acid washed coverslips (22 × 22 mm^2^), placed in 6 well plates containing nutrient broth and incubated for 72 hours at 37°C to form biofilm. After incubation, the non-adherent cells were removed by washing thrice with sterile 1XPBS. The biofilm formed on coverslip was then transferred to new 6 well plates. For phagocytosis assay, both live and heat inactivated biofilms were used.

For heat inactivation, two different conditions were used: heat treatment at 56°C for 30 minutes in water bath and 56°C for 1 h using bacteriological incubator. After heat treatment, both live and heat inactivated biofilms were stained with ConA-FITC (30 μg/mL) and propidium iodide (1 μg/mL) and observed using Fluorescent microscope.

### Phagocytosis assay

To study the phagocytic response of macrophage towards the biofilm, Raw264.7 macrophage cell line was used. The cells were plated at a density of 10^5^ cells/mL with complete DMEM medium without antibiotics and were then presented to the biofilm (FITC stained heat inactivated and live biofilm). The interaction study was performed by including the following groups: i. Non-activated macrophages presented to live biofilm, ii. non-activated macrophages presented to heat inactivated biofilm for 6 hours [to know the interactions between non-activated macrophage and biofilm], iii. activated macrophages presented to live biofilm, and iv. activated macrophages and heat inactivated biofilm.

The macrophages were activated by 3 *μg/mL* LPS (Sigma) and 100 pmol IFN-γ for 6 hrs. [In our previous study with planktonic cells, 4 hours incubation was used to activate macrophages by LPS. Since we presumed that biofilm interactions need more time, we extended the time to 6 hours (15)].

After incubation, the coverslips were washed with 1XPBS and treated with 0.005% trypan blue to quench out the ConA-FITC from biofilm cells [16] that were not engulfed by macrophages. After washing, macrophage cells were stained with Giemsa stain for 15 minutes and then observed under fluorescent microscope using a green filter for FITC stained biofilm, while stained macrophage were observed in bright field. The phagocytosis rate was calculated by counting,

Phagocytosis rate (%) = Macrophage with internalized biofilm – Macrophages without internalized biofilm/ Total number of macrophages X 100

### TLR2, iNOS and cytokine expression

To analyse the expression pattern of TLR2, iNOS and cytokine expression in Raw 264.7 macrophages during the phagocytosis of heat inactivated and live biofilm, the following experimental groups were included.

Group 1: Macrophages, Group 2: Activated macrophages, Group 3: Non-activated macrophages presented to live biofilm, Group 4: LPS activated macrophages presented to live biofilm, Group 5: Non-activated macrophages presented to heat inactivated biofilm, Group 6: LPS activated macrophages presented to heat inactivated biofilm.

After six hours of macrophage and biofilm interactions, the cells were washed with DEPC treated 1XPBS. Total RNA was then extracted from macrophages using the TRIzol method and converted into cDNA by PrimeScript™ RT-PCR Kit (Takara). The following genes were amplified, TLR2, iNOS, proinflammatory cytokines such as IL-β1, IFN-γ, IL-6, IL-12, IL-4 and TNF-α, anti-inflammatory cytokines IL-10 and GAPDH. These genes were amplified by initial denaturation at 95°C for 5 hours followed by 40 cycles at 95°C for 30 seconds. The different annealing conditions (30 seconds) are mentioned in the table and extension was at 72°C for 30 seconds. The PCR primers and amplification conditions are given in **Table 1.** The primer pairs were confirmed to amplify the genome DNA fragments of macrophage from control group (macrophages alone). All the groups were tested in triplicate on a real time PCR system (Eppendorf, Germany) using DyNAmo Flask STBR Green qPCR kit (Thermo Scientific) and analysed with the 2^−ΔΔCt^ method and normalized with GAPDH and control.

**Table 1.**
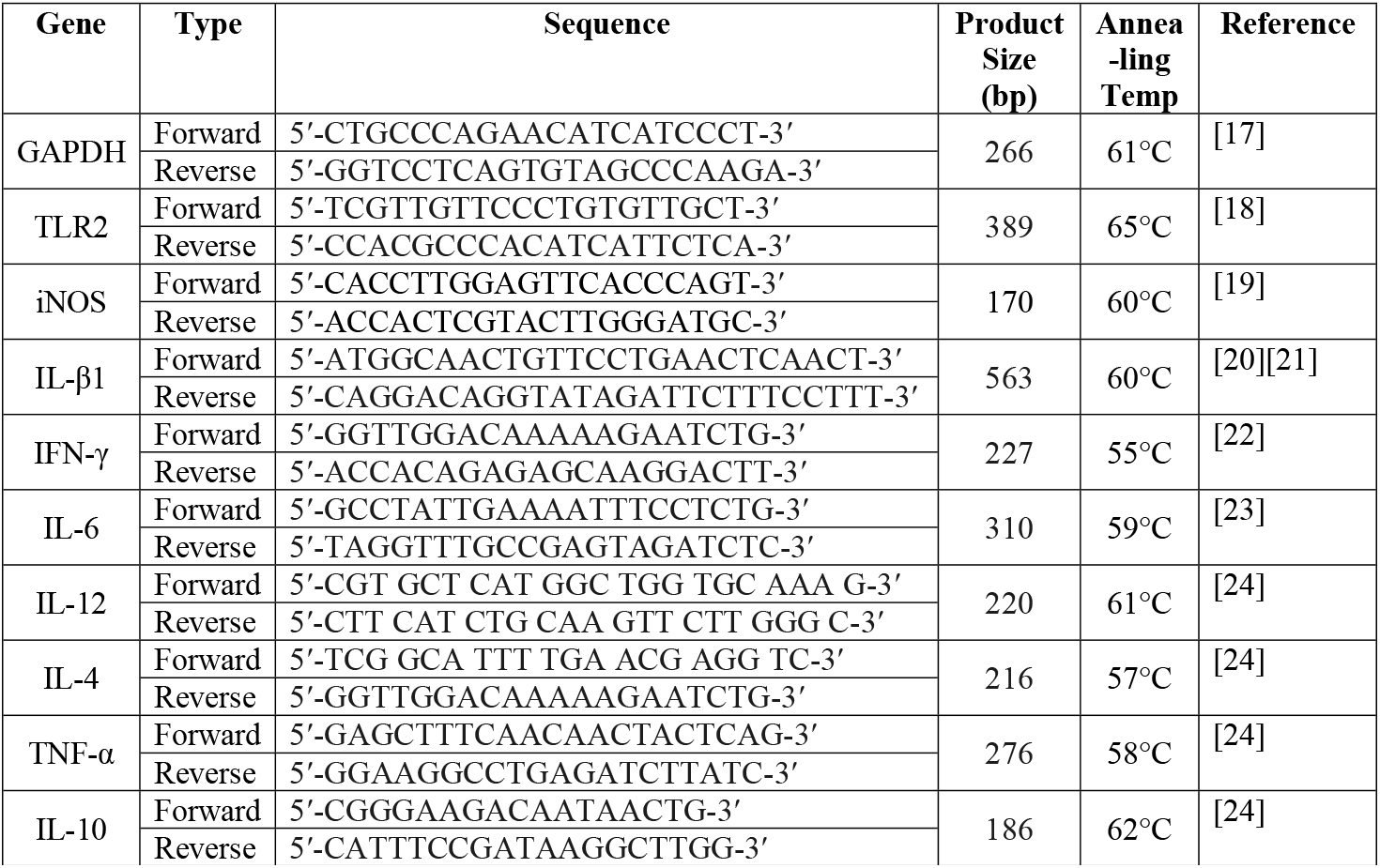
Primer sequence used for the cytokine gene expression

### Statistical analysis

All experiments were performed in duplicates and the data analysis was executed in GraphPad Prism 6. Two way ANOVA followed by post hoc test (Tukey’s multiple comparison) were performed to test statistical significance for multiple comparison. All graphs were prepared with GraphPad Prism 6 and were expressed as the mean ± standard deviation (SD) of triplicates.

## Results

### Heat treated *K. pneumoniae* biofilms

**Figure 1**, shows a visual comparison of heat-inactivated biofilms using water bath and incubator. The dead cells were observed in both the conditions. However, the biofilm cells are less in water bath treated when compared to incubator treated. The picture also reveals that the biofilm matrix was not disrupted during heat inactivation at 56°C for 1 h using bacteriological incubator. The untreated biofilm was shown to have dense live cells. Hence biofilms inactivated using incubator was selected for further study.

**Fig 1.**
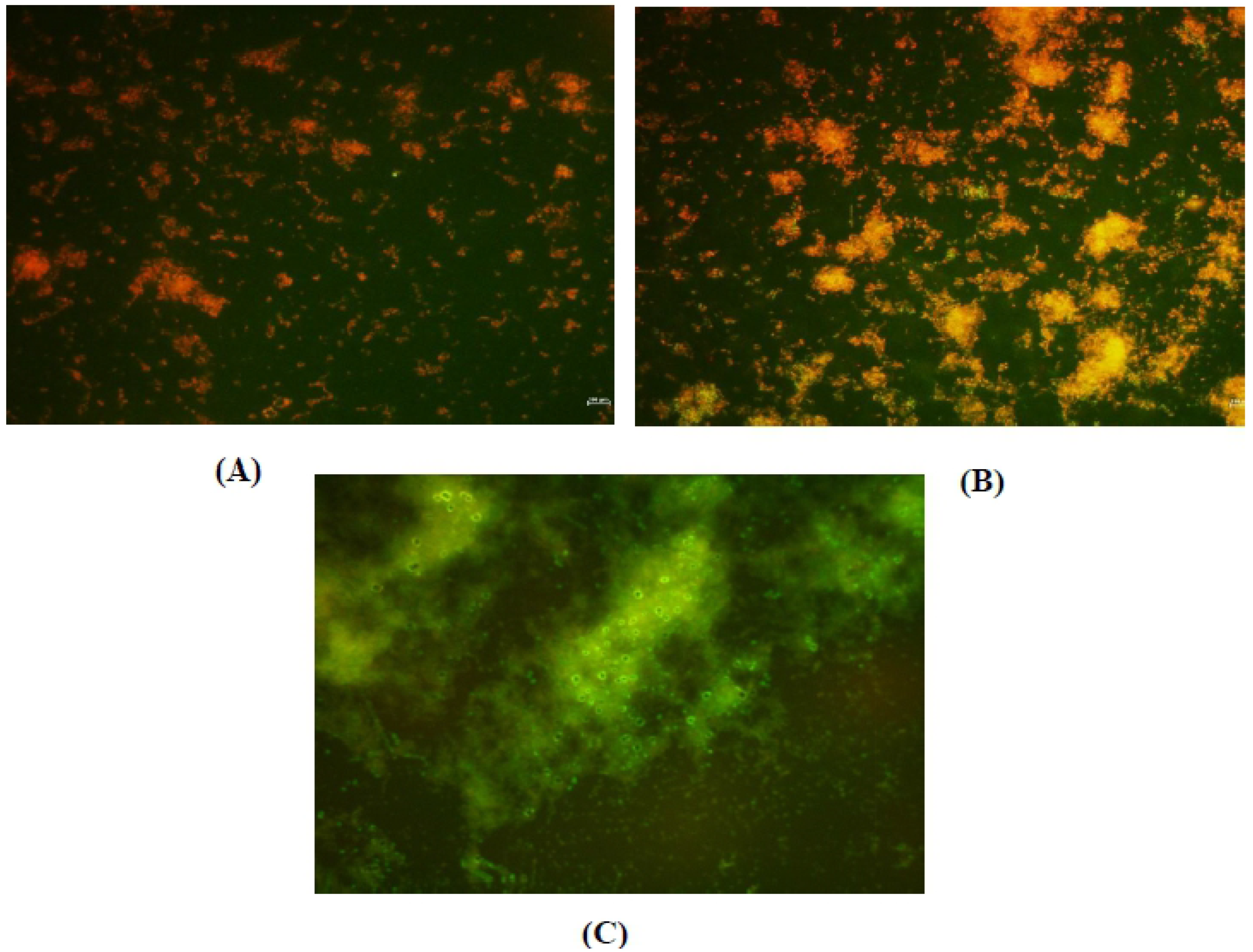
Heat-inactivated *K. pneumoniae* biofilm. **(A)** Heat treatment at 56°C for 30 minutes in water bath; **(B)** 56°C for 1 h using bacteriological incubator; (C) Untreated biofilm

### Phagocytic response of macrophage towards *K. pneumoniae* biofilms

To understand the role of macrophages in phagocytosis of *K. pneumoniae* biofilm, we co-incubated both these cells together under different treatment conditions as mentioned. LPS was used to stimulate the macrophages and both heat-inactivated or live biofilms were used for interaction. As can be seen from the results (**Fig 2**). Phagocytic rate of non-activated macrophage against heat inactivated biofilm was around 7±1.8% as compared to that against live biofilm (9.4±2%). This shows that there is certainly a basal response by macrophages against *K. pneumoniae* biofilms. However, when the macrophage were either pretreated or co-treated with LPS before exposure to *K. pneumoniae* heat inactivated or live biofilms, there was a significant increase in phagocytic rate. In the case of heat inactivated biofilm the phagocytic rate was 14.4±3.7 % and 15±2.8% for LPS pretreatment and co-treatment, respectively. A similar result was obtained when live biofilm was used (LPS pretreatment: 13± 3.3%; LPS co-treatment: 16±2.2%). It should be noted here that presence of LPS during macrophage interaction with either heat-inactivated or live biofilms consistently produced an increase in phagocytic response thought it was statistically insignificant (**S1 Table**).

**Fig 2.**
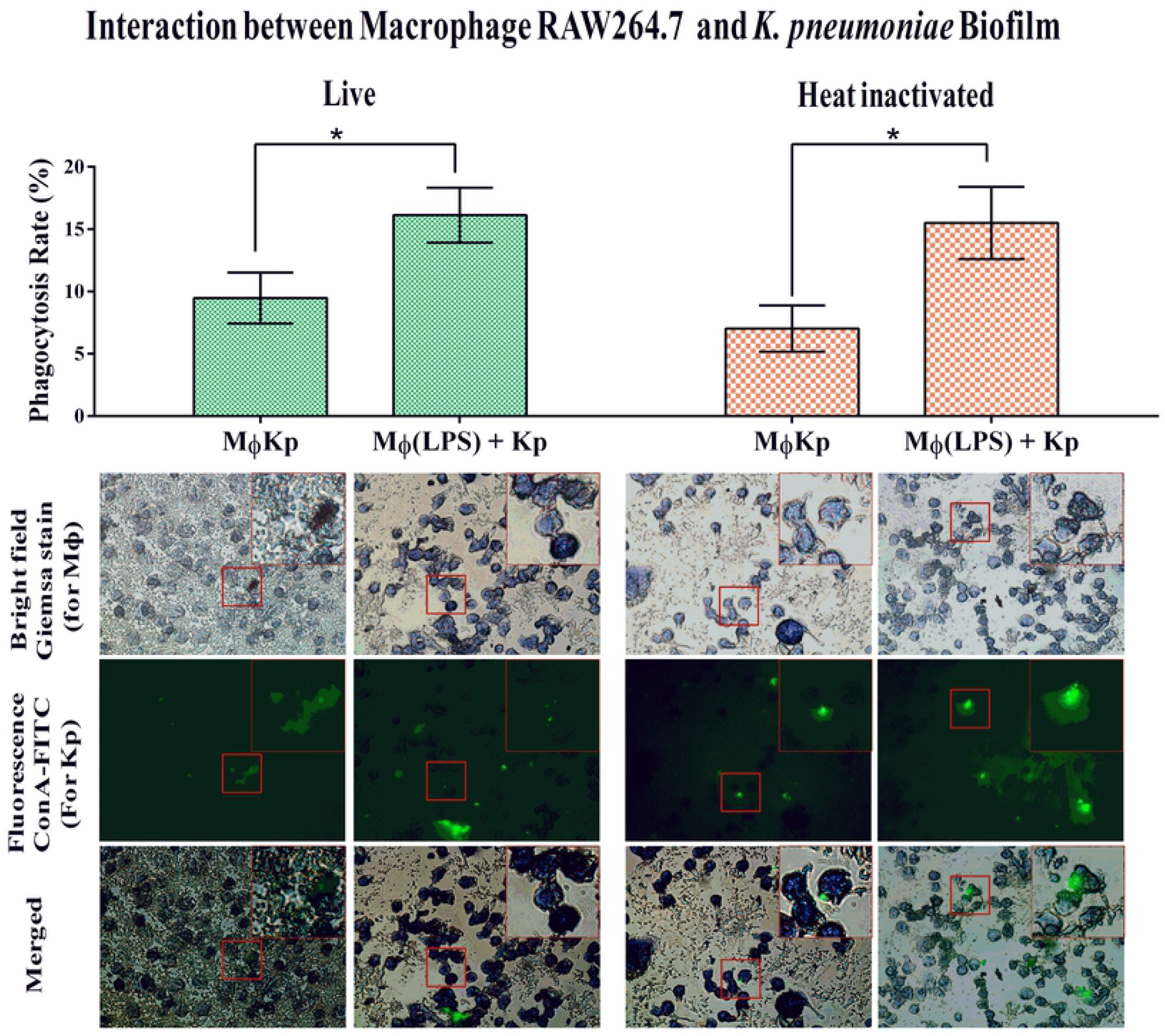
Interaction between Macrophage (MΦ) Raw264.7 and *K.pneumoniae* biofilm. (1) Non-activated MΦ exposed to the biofilm; (2) Phagocytosis assay in the presence of LPS and IFN-γ (co-treated). (A) Heat inactivated biofilm; (B) Live biofilm. Increased Phagocytic response was observed in both the cases of heat inactivated and live biofilm in the presence of LPS

### Cytokine responses of RAW 264.7 macrophage to *K. pneumoniae* biofilms

Another aspect of macrophage-microbial interaction is the production of cytokines by macrophages as a response to both recognition and phagocytic-killing of the microbe. To test whether this is true in the case of macrophage interaction with *K. pneumoniae* biofilms as well, we allowed LPS-activated macrophage to interact with either heat-inactivated or live biofilms of *K. pneumoniae* and analysed both pro- and anti-inflammatory cytokine gene expression together with TLR2 and iNOS genes. As can be observed from the results, there was a general increase in all the pro-inflammatory cytokines (IL-β1, TNF-α, IL-6, IL-12, IFN-γ), TLR2 and iNOS when LPS was used for macrophage activation (group 2), as is well known. However, use of LPS for activation resulted in a significant increase in anti-inflammatory IL-4 cytokine gene while there was a strong inhibition in the expression of IL-10 cytokine, when compared to unstimulated macrophages (group 1). Thus, it is clear that macrophage activation by LPS is a well-recognized prerequisite for setting up of pro-inflammatory immune responses. Using this as a background data, we next introduced both LPS-activated or inactivated macrophages to either heat inactivated or live biofilms. When comparing the results of both unactivated macrophage with either heat inactivated (group 3) or live biofilm (group 5) with that of group 2 cells, it is clear that there is no significant increase in any of the genes analysed, suggesting that LPS activation is very essential for inducing proinflammatory cytokine responses in macrophages. However, when group 3 responses are compared with that of group 5 cells there appears to be some change in gene expression. This is seen especially for TLR2 and IL-12 that showed a significant increase of 5-fold and 1.4-fold, respectively, for group 5 cells when compared to group 3 cells (group 3 IL-12 expression was in the negative). On the other hand group 5 macrophages showed a moderate increase in gene expression when compared to group 3 cells, while IL-10, as expected, remained unchanged (**Fig.3**).

**Fig 3.**
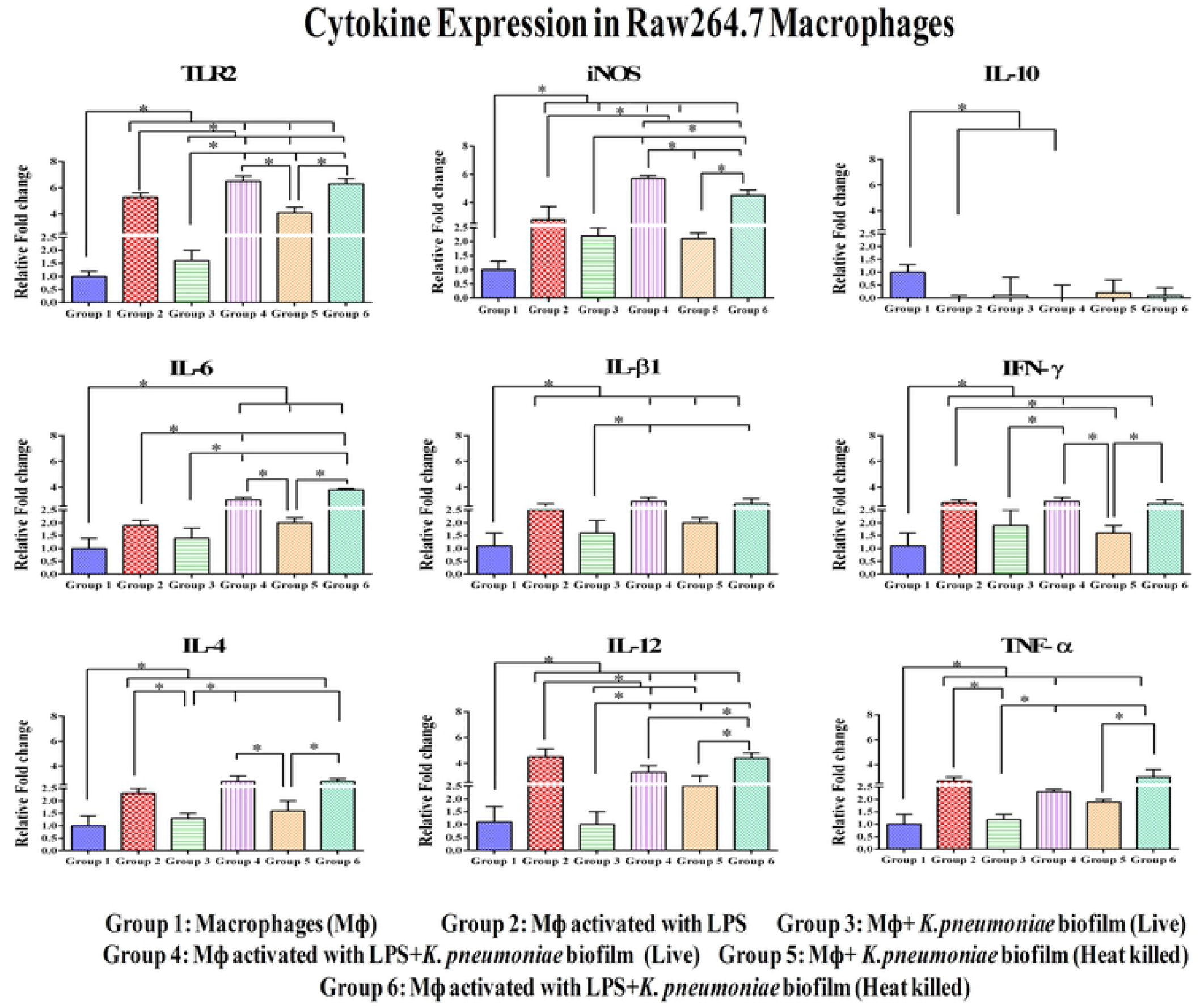
Cytokine gene expression assay. Evaluation of cytokines in RAW 264.7 Macrophage cell lines (MΦ) in various conditions. Heat inactivated or live biofilms of *K. pneumoniae* exposed to activated MΦ showed increased level of cytokines(TLR2, iNOS, IL-6, IL-β1, IFN-γ, IL-4, IL-12, TNF-α) except IL-10. Two way ANOVA followed by Tukey’s multiple comparison tests were performed. * represents significant fold increase (p < 0.05).

When macrophages were activated with LPS and then exposed to heat inactivated (group 6) or live (group 4) biofilms there was a significant increase in all the genes analysed, expect for IL-10 that showed no change. When comparing group 4 cells with group 3, it was observed that TNF-α showed the highest increase of 6-fold, followed by IL-6 and IL-4 (5-fold each), iNOS (4-fold), IL-β1 (3-fold) and IFN-γ (2-fold). A similar but more or less a consistent enhancement was observed across all the genes analysed in the case of group 6 cells when compared to group 5 macrophages, with increase ranging from 1.7 fold to 3-fold. These groups of cells showed a consistent increase in gene expression when compared to group 4 macrophages. Perhaps heat treatment of biofilm modifies certain surface properties producing these differences. Nevertheless, the highest response was observed for TLR2 in case of group 4 macrophages wherein it showed an increase of 9-fold over its control (group 3) macrophage. Taken together these results suggest that (1) LPS activation of macrophage is essential for better phagocytic response towards *K. pneumoniae* biofilm and, (2) activated macrophages are able to produce a stronger cytokine response when exposed to live biofilm, indicating better recognition of live biofilm cells by macrophages (**S2 Table**).

## Discussion

*K. pneumoniae* is a nosocomial pathogen whose resistance to antibiotics has become a problem worldwide and this is partly owed to its ability to form biofilms [25]. Such biofilm formation is seen quite commonly with MDR strains and extensively drug resistant *K. pneumoniae* [26]. Together with fimbriae production, biofilm production has become a major virulence mechanism [27] leading to immune evasion and establishment of infection [28]. Although there are studies to show the immune clearance of *K. pneumoniae* strains by the cells of the immune system [25], the basis for such recognition and interaction is still not fully understood, and moreover most studies have looked at only the planktonic or heat-inactivated cells. Thus, there is a general lack of information on the *K. pneumoniae* biofilm-innate immune cell interaction and this formed the basis of this study.

To understand the importance of biofilm in immune evasion, we used both heat-inactivated and live biofilms as targets and our results show that unactivated RAW 264.7 macrophages indeed show a basal phagocytic response towards biofilms, irrespective of the nature of the biofilm. However, we were able to obtain a higher phagocytic rate when macrophages were activated with LPS, before being exposed to biofilm. Even in this case the activated macrophages failed to distinguish between heat-inactivated and live biofilms and on an average the phagocytic rate was around 15% for either heat-inactivated or live biofilms. This suggests that RAW 264.7 macrophages have the ability to phagocytose biofilm strains of *K. pneumoniae* based on possible surface recognition. However, macrophage activation is needed to enhance this response, which suggests that the priming step is important for the activation of the innate immune response [29]. Nevertheless the rate of phagocytosis was an average 15% for both biofilms and this could be due to a variety of factors such as incubation time, role of biofilm in suppressing phagocytosis etc. [25]. Similarly previous *in vitro* experiments demonstrated minimal *S. aureus* phagocytosis [30]. In case of *S. epidermidis* biofilm, impaired phagocytosis and reduced activation of J774A.1 macrophages were noticed [31]. In our previous study we have shown the ability of the activated RAW 264.7 macrophages to phagocytose heat-inactivated planktonic *K. pneumoniae* clinical strain [15] by 32% and this difference we believe is due to the use of biofilm as a target here. Another criterion that could be considered for such an interaction is the exopolysaccharide and thus it remains to be seen if macrophage activation by exo-polysaccharide of *K. pneumoniae*.

Having shown the ability of activated macrophages to better phagocytose biofilms, we made an attempt to understand the ability of the biofilm to modulate cytokine, TLR2 and iNOS gene expression in the macrophages. It is already proven that the resistant strains of *K. pneumoniae* have the ability to not only resist phagocytic killing but also alter the polarisation state of the macrophages [32], which could be important in determining successful establishment of infection by *K. pneumoniae*. Pattern of cytokine secreted is one of the major mechanisms that determines macrophage polarisation [33] and microbes especially resistant ones are well known to skew that innate immune responses to a more anti-inflammatory type [34]. Our study shows that when activated macrophages were exposed to heat-inactivated or live biofilms, there was a significant increase in pro-inflammatory cytokine genes together with expected increase in TLR2 and iNOS. Interestingly, anti-inflammatory IL-10 showed no upregulation in any of the treatment groups. Surprisingly, both heat-inactivated and live biofilms induced similar upregulation of pro-inflammatory genes in macrophages, suggesting a minor role of exo-polysaccharide in modulating macrophage cytokine responses. However, we would like to point out that this needs more detailed study using isolated exo-polysaccharide. At this juncture we would like to point out that in the case of group 3 macrophages (unactivated + exposed to live biofilm) and group 5 macrophages (unactivated + exposed to heat-inactivated biofilm) the gene expression was lower than group 2 (activated macrophage alone) macrophages, suggesting the possibility of (1) suppression of macrophage immune responses by the biofilm, and (2) the essentiality of macrophage priming by LPS + IFN-γ, at least in vitro. Nevertheless the results from gene expression analyses clearly show that during RAW 264.7 macrophage interaction with *K. pneumoniae* biofilm, there is a modulation of the macrophage responses towards a pro-inflammatory one and this could be important in increasing the clearance efficacy of innate immune cells. Nevertheless, this modulation was not similar for LPS activated Raw264.7 macrophages and planktonic *K. pneumoniae* cells [15]. The cytokine expression was found to have a significant increase in, IL-4 (8-fold), IL-12 (5-fold), TNF-α (7-fold), IFN-γ (17-fold), whereas in case of biofilm, the cytokine modulation was very minimal ranging between 1-3 fold increase in cytokine expression. Taken together, the *in vitro* results suggest that *K. pneumoniae* in biofilm mode elicits minimal phagocytic response and cytokine expression by macrophages.

## Conclusion

The *in vitro* model was developed to culture *Kebsiella* biofilm and macrophage to mimic *in vivo* phagocytic response of macrophage towards biofilm. The *in vitro* results suggest that *K. pneumoniae* in biofilm mode has minimal phagocytic response and cytokine expression by macrophages. Also, these results show that for a better *K. pneumoniae* biofilm recognition and clearance by macrophages, their activation state is essential in determining both the phagocytic rate and polarisation state, under *in vitro* conditions. Perhaps this could be important in terms of devising strategies for treating extremely resistant biofilm producing strains of *K. pneumoniae*.

## Acknowledgment

We thank Science and Engineering Research Board (SERB), Department of Science and Technology, New Delhi for funding. We thank SASTRA Deemed University for providing the research facilities and infrastructure.

## Funding

Science and Engineering Research Board (SERB), Department of Science and Technology, New Delhi (EMR/2016/007613) to JP for funding this research and DST-FIST funding (No: SR/FST/ETI-331/2013) for fluorescence microscope support.

## Supporting information

**S1 Table. Statistical analysis of macrophage interactions.** Two way ANOVA followed by Tukey’s multiple comparison tests were performed for the macrophage interactions exposed to the heat-inactivated or live biofilms

**S2 Table. Statistical analysis of cytokine gene expression in Raw264.7 macrophages.** Two way ANOVA followed by Tukey’s multiple comparison tests were performed for the cytokine gene expressions (TLR2, iNOS, IL-6, IL-β1, IFN-γ, IL-4, IL-12, TNF-α and IL-10)

